# Column-Like Subnetwork Reconstruction in Motor Cortex from Graph-Based 3D High-Density Two-Photon Calcium Imaging

**DOI:** 10.1101/2025.06.17.660119

**Authors:** P. Aymard, J-C. Boffi, H. Asari, R. Prevedel, D. Holcman

## Abstract

How precise 3D interactions among cortical neurons underlie layer-specific computations remains elusive. We developed a graph-framework to infer functional connectivity from fast volumetric two-photon Ca^2+^ imaging in the awake mouse primary motor cortex. By converting deconvolved traces into binary spike trains, removing population bursts, and applying an adaptive, layer-specific statistical threshold, we reconstructed a directed, weighted network of ∼1,000 neurons. Decomposition into strongly connected components revealed ∼10-cell subnetworks, predominantly in layer II/III but often bridging to layer Va. Across six 20-min recordings, we found that (1) layer II/III dominates connectivity, (2) feedback (Va→II/III) links are more numerous and stronger than the feed-forward (II/III→Va) ones, and (3) information flows in ≤ 6 synapses. We uncovered seven geometrical and dynamical motifs—ranging from compact columns to elongated diagonals—each with characteristic event sizes and durations. These findings reveal diverse, column-like microcircuits in M1 with a net ascending flow, suggesting that such subnetworks form elemental processing modules for motor control.

## Introduction

Although cortical columns have long been proposed as the neocortex’s elementary processing units [1], how their three-dimensional (3D) functional pathways support complex computations remains poorly understood. Unraveling this micro-scale architecture is essential for deciphering key processes such as perception, cognition, and refined motor control [2, 3, 4]. In sensory areas, sequential two-photon imaging has provided indirect evidence of radial organization through receptive-field mapping [5, 6, 7], but such approaches cannot capture simultaneous activity across layers and thus fall short of revealing true functional connectivity.

Recent advances in volumetric *in vivo* imaging—most notably scanned temporal-focusing two-photon (sTeFo 2P) microscopy—now permit near–simultaneous recordings of thousands of neurons across hundreds of micrometers in depth [8, 9, 10, 11]. This dense, high-speed sampling opens the door to reconstructing three-dimensional networks, yet a systematic, data-driven framework for inferring layer-to-layer connectivity from such recordings has not been established. Prior efforts to infer functional networks from calcium imaging [12] either relied on simulated data, two-dimensional analyses, or neglected cortical layer diversity and 3D topology [13, 14, 15, 16, 17, 18, 19, 20, 21, 22, 23].

Here, we introduce a scalable graph-pipeline that transforms volumetric calcium time series into directed, weighted networks of cortical neurons that mirror neuronal networks. By converting deconvolved traces into binary spike trains, removing population bursts via a background-matching filter, and applying adaptive, layer-specific statistical thresholds, we infer high-confidence functional links between individual cells. Decomposition into strongly connected components (SCCs) reveals ∼10-cell subnetworks that predominantly form column-like motifs in layer II/III and often bridge to layer Va. Across six 20-min recordings in the primary motor cortex (M1) of awake mice, we identified seven distinct SCC archetypes, demonstrate a net feedback (Va→II/III) information flow in ≤ 6 synaptic steps, and uncover highly connected hub neurons that reinforce local dynamics. Our approach both confirms longstanding hypotheses about cortical columns [1, 24, 25] and unveils previously unrecognized microcircuit motifs—providing a generalizable toolkit for mapping functional architecture throughout the neocortex.

## 1 Results

To unveil the fine-scale organization of functional subnetworks in M1, we start by developing a computational pipeline that converts volumetric calcium signals into graph representations of neuronal connectivity. Below, we summarize the key steps—from raw fluorescence traces to a directed, weighted network.

### Volumetric two-photon imaging of large-scale mouse motor cortex activity in awake mice

We performed fast volumetric calcium imaging using sTeFo 2P microscopy [8] in awake, head-fixed C57BL/6J mice. We recorded one session on 6 different animals resulting in 6 datasets of ∼1,000 neurons within a 470×470×450 *µ* m^3^ volume at 4Hz. To target the M1 region controlling face movements—and avoid intermingled motor areas—we centered our field of view on the snout-associated zone [26]. Because mouse M1 is agranular [27], we classified somata deeper than 400 µm as layer Va and those above 400 µm as layer II/III. Raw ΔF/F_0_ time series were first motion- and neuropil-corrected, baseline-drift–adjusted, and denoised using the CaImAn toolbox [28]. We then deconvolved each trace and extracted spike times by identifying peaks with a prominence threshold of *p* = 0.015 (Fig. 1A). Although mice remained passive during imaging, three of six datasets exhibited transient, high-amplitude population bursts. To prevent these episodes from biasing our connectivity estimates, we applied a background-matching filter that selectively down-samples burst-associated spikes while preserving overall event continuity (see Methods 3.4.2 and Fig. 1B).

**Figure 1:**
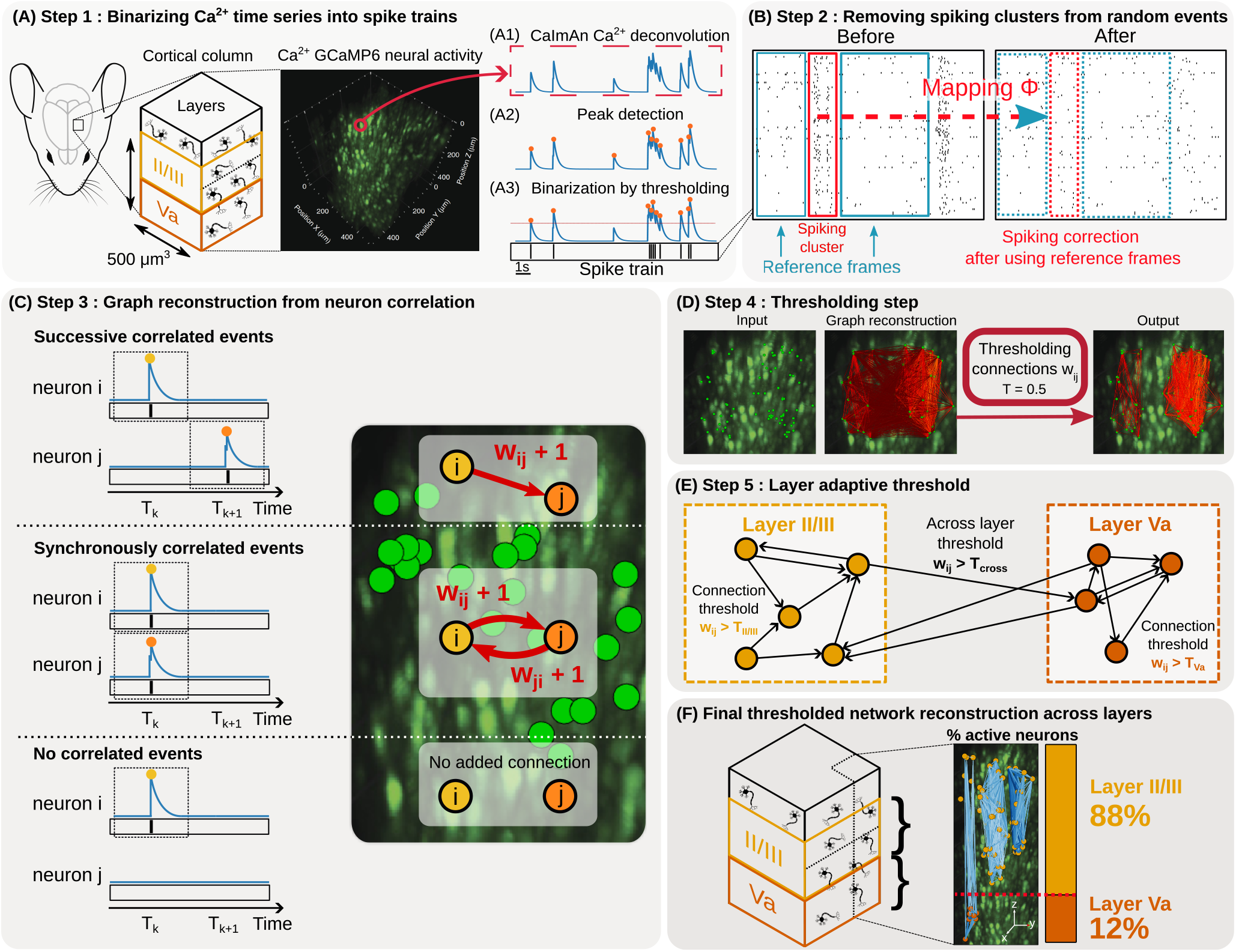
Network reconstruction. **A**. Ca^2+^ signals recorded from a 470×470×450 *µm*^3^ = 0.1 *mm*^3^ volume across layer II/III and Va from a mouse brain (left) Step 1: A1. GCaMP6f Calcium time series deconvoluted with CaImAn [28]. A2. Peak detection with prominence *p* = 0.015 (see Table 3). A3. Binarization of calcium events into spike trains using threshold *T*_*f*_ = 0.1. **B**. Step 2: Removal of spiking cluster events using function *ϕ* (see Methods SS2) that matches bursting firing rates to the background activity (blue). **C**. Step 3: Computing the matrix weight *w*_*i,j*_ which represent temporal correlation between two firing neurons *i, j* : a successful correlation between neuron *i* and *j* is associated with a +1 increment of *w*_*i,j*_. For a simultaneous correlation event, both *w*_*i,j*_ and *w*_*j,i*_ are incremented by one. When no correlation occurs, the weight remains unchanged. **D**. Step 4: Thresholding procedure to remove coincidental connections between neuron *i* and *j* when the weight *w*_*i,j*_ *< T*. **E**. Step 5: Thresholding procedure is adapted to each layer population (see Method 3.4.5). **F**. Final output: Network reconstruction of cortical layers. Layer repartition : 88% for layer II/III vs 12% for layer Va of active neurons with at least one connection.

### Reconstructing 3D functional connectivity across layers

To infer layer-to-layer functional links despite our relastively low 4Hz sampling rate and 20min recordings, we adopted a recurrence-based, statistical correlation framework. We computed pairwise weights by counting both synchronous and consecutive spike events between every neuron pair (Fig. 1C), reasoning that true functional connections will reappear reliably over long recordings. By aggregating these co-activation counts across the entire session, we obtained a directed, weighted adjacency matrix in which stronger edges reflect more consistently recurring interactions. This approach ensures that only pathways with repeated temporal associations—rather than isolated coincidences—survive into the reconstructed 3D network.

To prune spurious co-activations, we derived an adaptive, layer-specific threshold from a data-driven null model. In this model, independent Poisson firing (see Method 3.4.5)—parameterized by each layer’s empirical rate distribution and the 20min recording length— yields the maximum coincidental co-activation count expected by chance (Fig. 1D–E). Only edges whose weights exceed this confidence bound (p*<*0.05) are retained, ensuring that our 3D network reflects repeatedly recurring interactions rather than isolated coincidences.

Applying this criterion across six datasets, we reconstructed functional graphs encompassing on average 33.9 ± 12.1% of imaged neurons. This conservative fraction is likely due to both outside-volume connectivity and the stringent confidence threshold. Within this subnetwork, 88% of retained neurons reside in layer II/III versus 12% in layer Va—slightly enriched relative to their raw proportions (84% versus 16%), consistent with higher firing rates and stronger recurrent dynamics in the superficial layers (Fig. 1F).

### Quantitative connectivity patterns in layers II/III and Va

We represent the reconstructed network as a directed, weighted graph with adjacency matrix *A* ∈ ℝ^*N* ×*N*^, where *N* is the number of neurons (Fig. 2A). Each entry *A*_*ij*_ = *w*_*ij*_ counts the number of synchronous or successive co-activations between neuron *i* and neuron *j* (Fig. 1C); absent connections have *w*_*ij*_ = 0. After applying our statistical threshold (see Methods 3.6), we computed standard graph metrics.

**Figure 2:**
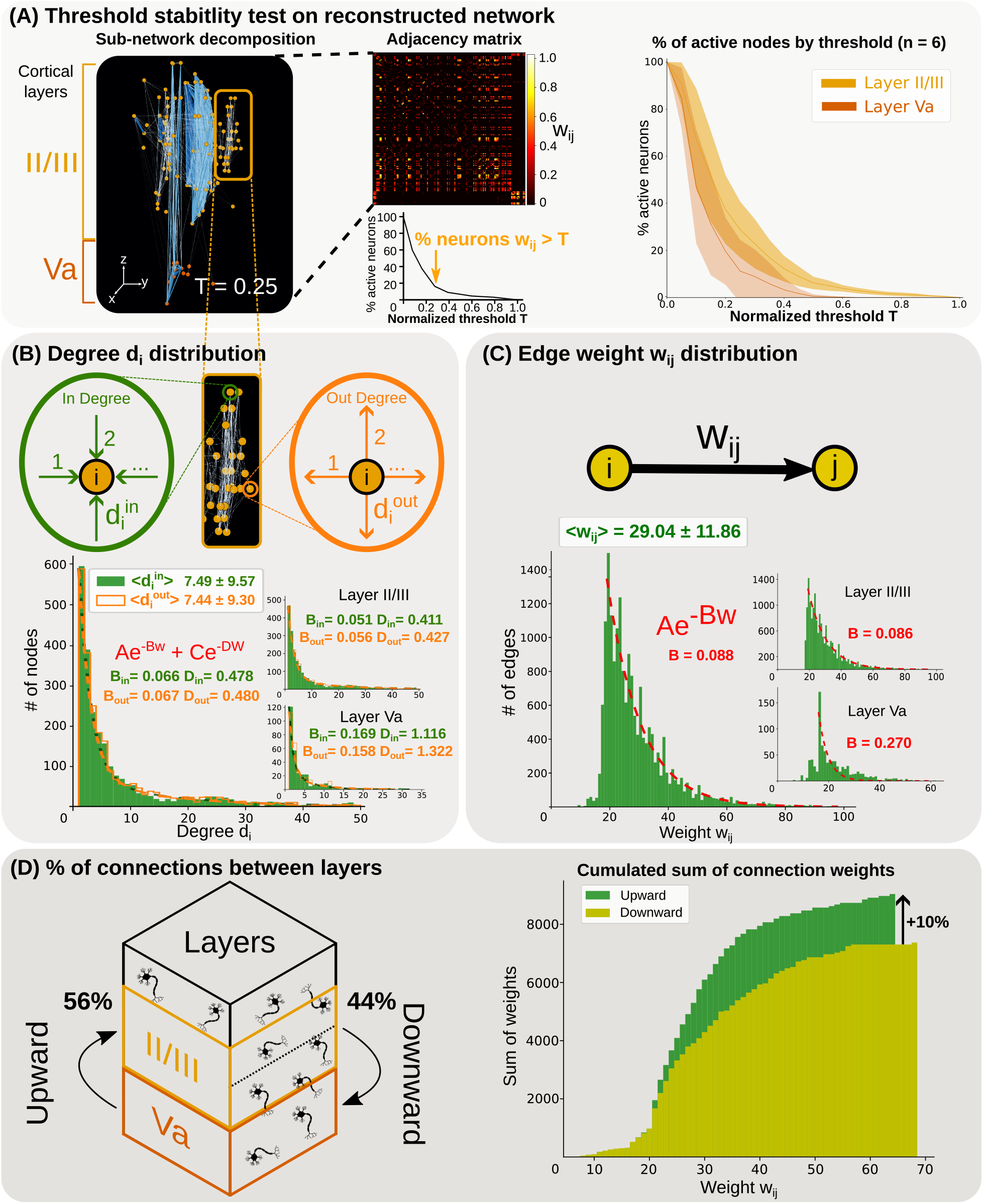
Structure and statistics of the neuronal network across layers II/III and Va. **A**. Left: example of a network reconstruction with threshold *T* = 0.25. Middle: normalized adjacency matrix *w*_*i,j*_*/max*(*w*_*i,j*_). Right: evolution of the proportion of active neurons with at least one connection between Layer II/III and Layer Va when increasing threshold *T* over the 6 experiments (6 mice). **B**. Upper : In/out degree definition. Lower: In (green) and out (orange) degree distributions over the entire layers and within layer II/III and Va. A sum of two decaying exponential is fitted over each degree distributio6n. **C**. Edge weight *w*_*i,j*_ distribution fitted by a single exponential (red). **D**. Left: Proportion of upward (resp. downward) connections from Layer Va to Layer II/III (resp. II/III to Va) in the network. Right: Cumulated sum of upward connections (green) and downward connections (yellow) weights.

The inand out-degree distributions exhibit a monotonic decay but feature a small “bump” in the in-degree between 18 and 30 connections, shifting its mean (7.44±9.30) slightly above the out-degree mean (7.49±9.57) (Fig. 2B). Layer-specific breakdown shows this excess indegree is driven by neurons in layer II/III, consistent with locally clustered activity. We fitted both degree distributions with a double-exponential model (*y* = *Ae*^−*Bx*^ + *Ce*^−*Dx*^), which outperformed single-exponential and power-law alternatives, suggesting two activation regimes: a background Poisson-like firing and a reinforced, hub-driven regime in layer II/III. This observation aligns with recent evidence that cortical networks rarely follow strict scalefree laws [29].

Edge weights—normalized by each dataset’s maximum—also follow an exponential decay (*y* = *Ae*^−*Bx*^), with a mean weight of 29.0±11.9 across layers (Fig. 2C). Again, layer II/III dominates the high-weight tail. Together, these results underscore the prominent role of superficial layers in structuring both the density and strength of functional links in M1.

Another critical insight emerged when we segregated inter-layer edges by their directionality. We found that when we pooled all connections between layer Va and II/III 56% of were ascending (Va→II/III), versus 44% descending (II/III→Va) (Fig. 2D, left). This result seems to be heterogeneous across dataset giving a less significant averaged result (50.1±13%). However, ascending links carried 10% more total weight than descending ones (Fig. 2D, right), an imbalance driven primarily by edges of intermediate strength (weights 20–40). These results reveal a robust feedback bias in functional information flow, with mid-range pathways disproportionately reinforcing the upward projection.

### Identification of high-density functional modules via strongly connected components

To identify densely interconnected sub-networks, we decomposed the directed graph into strongly connected components (SCCs), each defined by the property that every neuron is reachable from every other (Fig. 3A). Across six recordings, we identified on average 29.2 SCCs per dataset (≈175 in total), with a mean size of 9.8±9.2 neurons (Fig. 4A). Most SCCs formed compact, column-like clusters, though a minority appeared as multi-column outliers. We used DBSCAN [30] on SCC size and planar spread to isolate these outliers and then split them into elementary columns using a distance threshold *T*_*D*_ (Methods 3.5.1). After discarding trivial SCCs (fewer than *T*_*scc*_ neurons), 85% of modules localized entirely to layer II/III, 7% to layer Va, and 8% spanned both layers (Fig. 3B).

**Figure 3:**
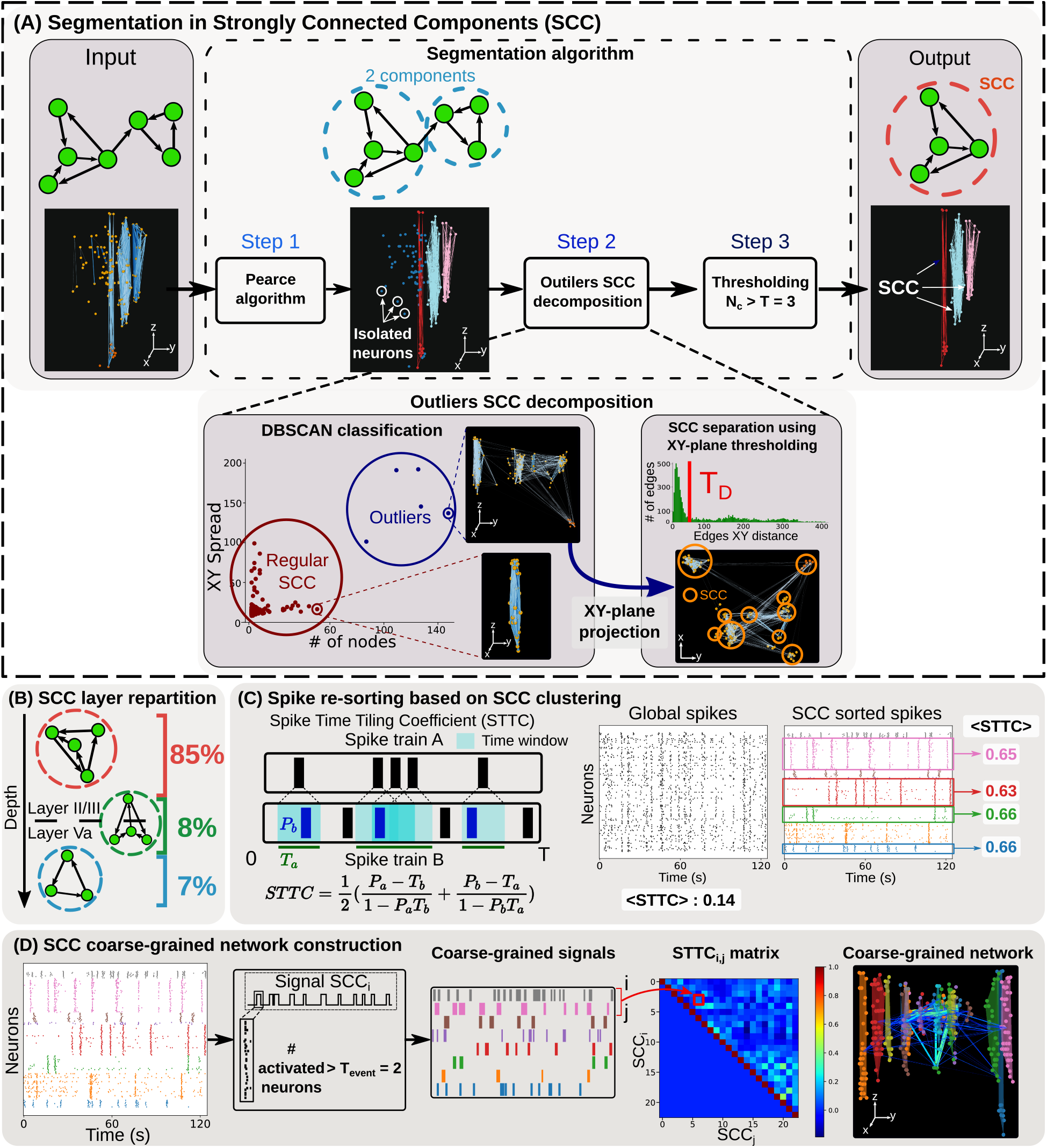
Decomposition of cortical layer network into sub-components. **A**. Segmentation of the network in strongly connected components (SCC) in three steps. Step 1: Pearce algorithm to decompose the network in SCC. Step 2: Outlier detection using the DBSCAN algorithm based on a planar projection (see inset) (see Method 8). Outliers are further decomposed into smaller SCC by removing edges with planar distance *d*_*XY*_ *< T*_*D*_ = 45 (see Table 3). Step 3: Constructing the final set of sub-networks by keeping the components where the number of neurons *N*_*c*_ *> T* = 3. **B**. Distribution of sub-networks across layers : 85% in layer II/III, 8% across layers, and 5% in layer Va. **C**. Left : illustration of Spike Time Tiling Coefficient (STTC) [31]. Middle : average STTC for all spike trains. Right : Spike trains sorted by sub-networks using sub-figure A and their associated STTC (pink, red, green, blue). **D**. Network coarse-graining pipeline. From left to right: Spike trains sorted by SCC, binarization of activation event for a number of consecutive activated neurons *> T*_*event*_ = 2, binarization results leading to coarse-grained representation of time events, STTC matrix associated to coarsed-grained netw7ork reduction and final 3D network decomposed into SCC.

**Figure 4:**
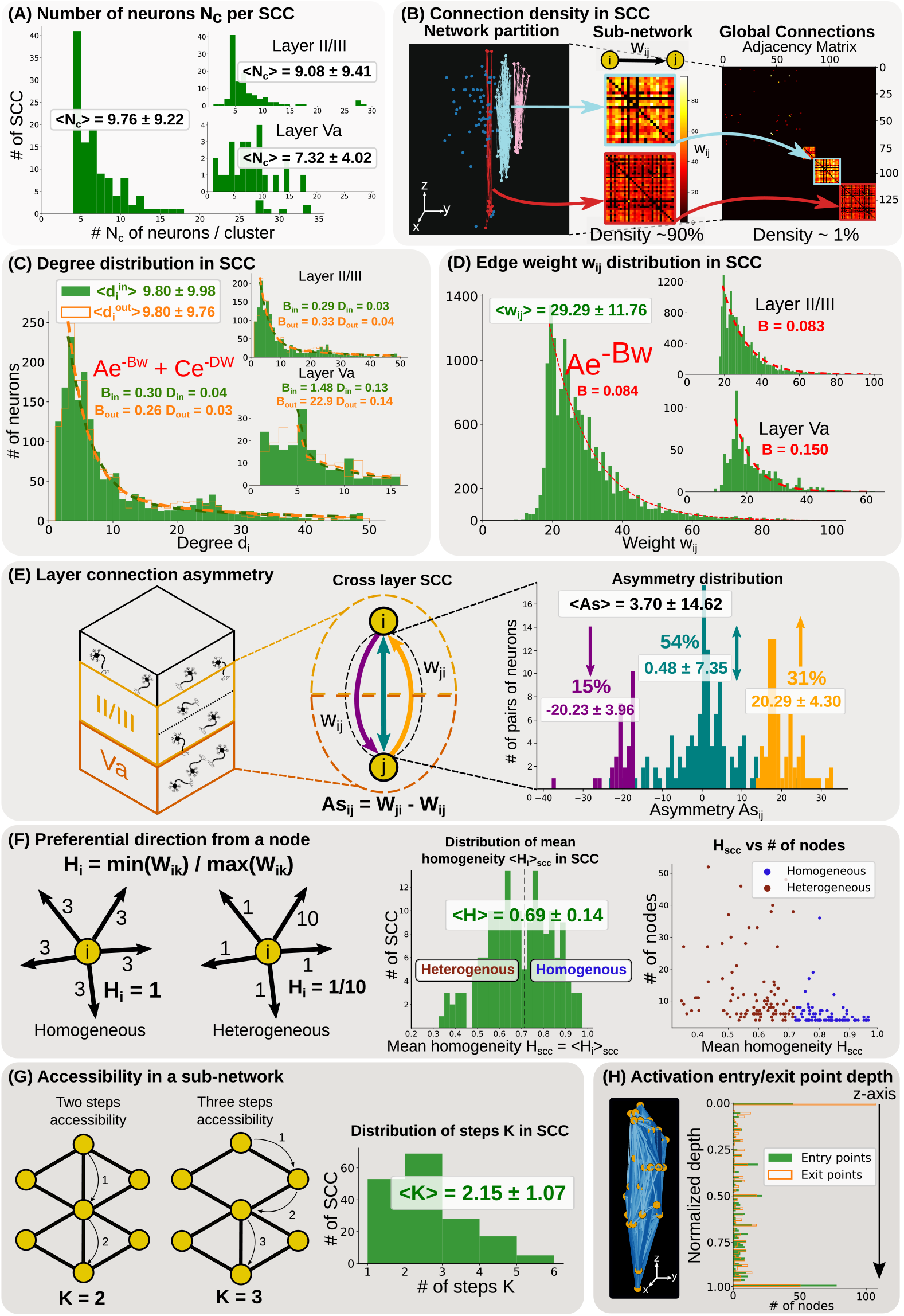
Structural and connectivity flow in cortical sub-networks. **A**. Distribution of SCC’s number of neurons *N*_*c*_ across layers. **B**. High density adjacency matrices of two SCCs and adjacency matrix of the entire network sorted by sub-networks. **C**. In (green) and out (orange) degree distributions for all SCCs (*n* = 175) over the entire layers and within layer II/III and Va. A sum of two exponential is fitted over each degree distribution.**D**. Edge weight *w*_*i,j*_ distribution for all SCCs fitted by a single exponential (red). **E**. Distribution of the asymmetry weight *As*_*i,j*_ between pairs of neurons of different layers within SCCs. **F**. Left : Statistics of preferential direction in neurons characterized by the homogeneity *H*_*i*_ = *min*(*w*_*i,k*_)*/max*(*w*_*i,k*_) coefficient. Middle : Distribution of the average homogeneity *H*_*i*_ withi9n SCCs, separated into two populations at threshold *H*_*scc*_ = 0.7 described either as more heterogeneous or homogeneous. Right : Homogeneity vs heterogeneity with respect to the number of nodes in SCCs. **G**. Left : Minimal number of steps *K* to cross a graph: two examples. Right: distribution of *K* within SCCs. **H**. Distribution of

Compared to the full network (edge density 0.010±0.002), SCCs exhibited dramatically higher connectivity (mean density 0.725±0.188), i.e., ∼70-fold increase in local connection density (Fig. 4B). Within SCCs, both inand out-degree distributions peaked at three connections—versus one in the global graph—reflecting the removal of dead-end nodes (Fig. 4C). Although the average degree remained comparable, SCCs preserved the “bump” in in-degree at 18–30 links, confirming that the hub-driven regime arises from these tightly layer II/III modules. Edge-weight distributions within SCCs (Fig. 4D) mirrored those of the overall network, indicating that our partitioning retains the full spectrum of functional link strengths.

### Directional flow and accessibility within subnetworks

Focusing on SCCs that span both layers, we quantified inter-layer bias using an *asymmetry* index (Methods 3.6), defined as *A*_*ij*_ − *A*_*ji*_ for each cross-layer pair. Positive values denote net Va→II/III flow, negative values II/III→Va. Across all cross-layer SCCs, the asymmetry distribution is significantly shifted toward positive values (Fig. 4E), confirming a predominant feedback information bias from layer Va to II/III within M1. Bidirectional edges—where *A*_*ij*_ ≈ *A*_*ji*_—form a synchronously activating backbone that does not contribute to net flux. To assess local wiring uniformity, we introduced a *homogeneity* metric, the ratio of a node’s smallest to largest outgoing weight. SCCs bifurcate into two regimes: “homogeneous” modules with evenly balanced links, and “heterogeneous” ones with pronounced hub-to-periphery pathways (Fig. 4F). Small SCCs (*<*8 neurons) are equally split between these types, whereas larger SCCs (*>*8 neurons) are predominantly heterogeneous, suggesting that extended modules develop stronger preferential routes over longer activation windows.

Finally, by normalizing *A* to a transition matrix *P* and computing the smallest *k* such that all entries of *P* ^*k*^ are positive [32], we estimate the maximal number of synaptic relays required for full spiking redistribution. The distribution of *k* peaks at ∼2.2 (Fig. 4G), with an upper bound at six, and thus much fewer intermediate neurons suffice to bridge any two nodes across network layers. Consistent across SCCs, event-entry neurons are disproportionately found in deeper (Va) positions, while exit neurons cluster superficially (II/III) (Fig. 4H), mirroring the net upward flow.

### Diverse column structural organization revealed by unsupervised clustering

To catalog recurrent subnetwork archetypes, we applied an unsupervised classification to SCC feature vectors (Fig. 5A; Methods 3.5.3). Initially, SCCs were grouped by laminar extent—Va-only, II/III-only, or cross-layer—revealing that 85% reside in II/III (Fig. 3B). Focusing on the overrepresented II/III components, we extracted 27 geometric and functional metrics (e.g., component size, spatial spread, link density, homogeneity, spectral gap) and reduced them via a PCA analysis. Then we applied a K-means clustering (elbow method, K = 5) and partitioned these into five II/III-specific classes, which together with the two laminar classes yielded seven distinct subnetwork types (Fig. 5B; Table 1).

**Table 1:** Table of sub-networks subtypes characteristics.

|  | Type 1 | Type 2 | Type 3 | Type 4 | Type 5 | Type 6 | Type 7 |
| --- | --- | --- | --- | --- | --- | --- | --- |
| Name | Layer Va | Cross Layer | Outliers | Large | Short | Elongated | Small |
| Layer | Va | Both | II/III | II/III | II/III | II/III | II/III |
| # neurons $N_c$ | $\sim 7$ | $\sim 12$ | $\sim 8$ | $\gg 8$ | $\sim 8$ | $\sim 8$ | $\ll 8$ |
| Spatial organization | Short | Elongated | Random | Elongated | Short | Elongated | Short |
| Frequency % | 6.3 | 8.0 | 3.4 | 9.7 | 12.0 | 29.7 | 30.9 |

**Figure 5:**
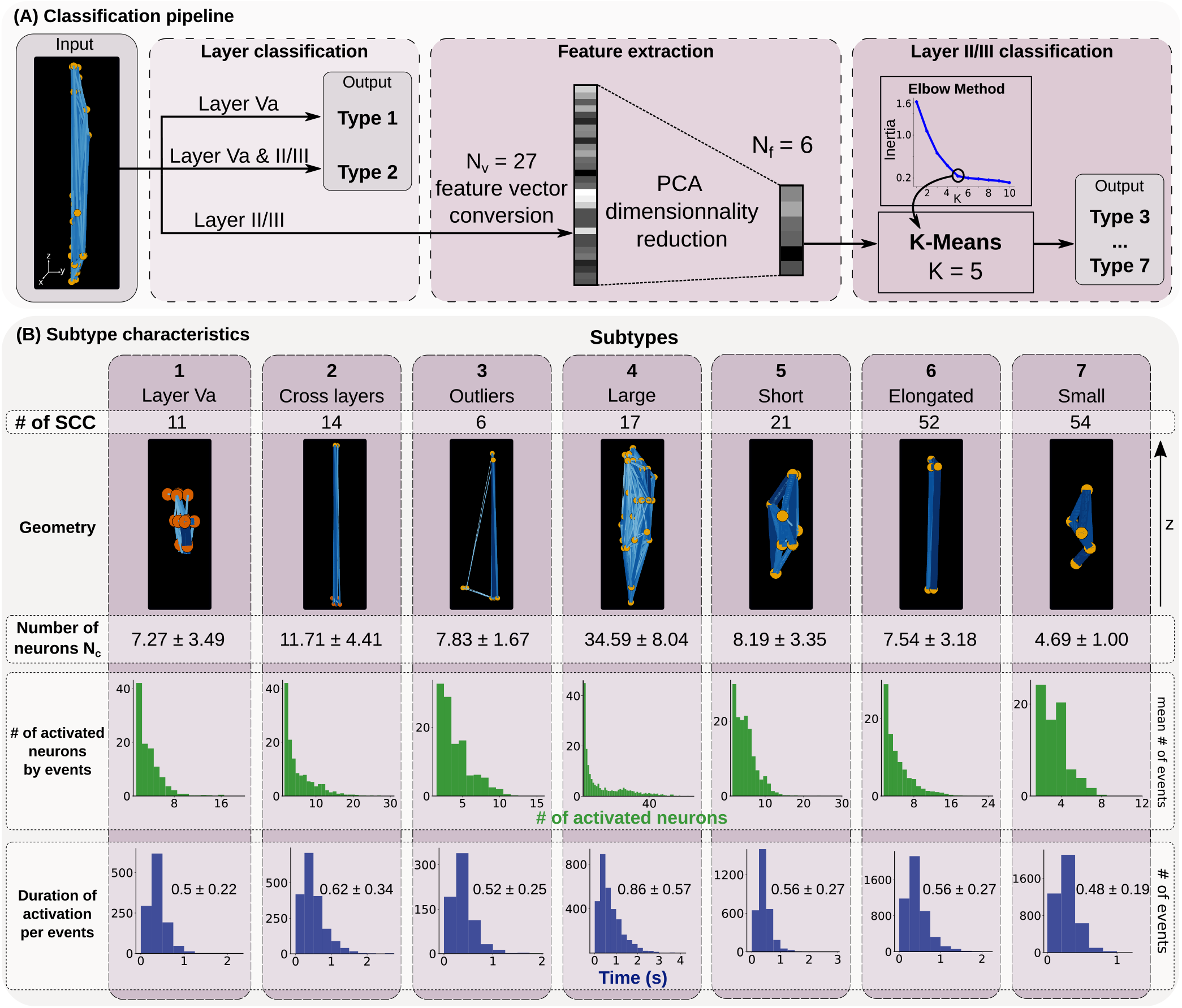
Subtypes network classification. **A**. Classification pipeline of sub-networks in 7 types: Step 1: Classifying sub-networks according to their layers. Step 2: Extraction of 27 features (see Table 2 and Methods) for sub-networks in layer II/III. Dimension reduction to dimension 6 using PCA. Step 3: Clustering of layer II/III sub-networks using K-Means algorithm parametrized with an elbow method (*K* = 5). **B**. Sub-network characteristics for the 7 subtypes: number of sub-networks in each class, their characteristic geometry, number of neurons *N*_*c*_, distribution of activated neurons per events and duration distribution (see Table 1 for more details).

We next analyzed activation “events,” defined as sequences of ≥2 SCC neuron firings that end when no further spikes occur. Notably, type 4 SCCs alone produced events recruiting ∼20–30 neurons synchronously spiking—precisely accounting for the in-degree “bump” seen earlier. This behavior leads to a highly correlated sub-networks where every neuron is strongly correlated. Other classes segregated into “short” motifs (compact, brief activations) and “elongated” motifs (longer, vertically extended sequences). Remarkably, when we normalized event size and duration by SCC neuron count, all seven types collapsed onto a common distribution, and both mean event size and duration scaled linearly with module size (Fig. 5B). These results suggest that—under passive conditions—the temporal dynamics of cortical microcircuits depend primarily on subnetwork size rather than laminar identity.

**Table 2:** Features extracted from strongly connected components.

| Feature type | Name | Symbol |
| --- | --- | --- |
| Geometrical | Number of neurons | $N_c$ |
| | Maximal distance and vector between neurons | $d_{max}$ |
| | Minimal distance and vector between neurons | $d_{min}$ |
| | Angle between the maximal and minimal vector between neurons | $\theta$ |
| | Center of mass | $m_1$ |
| | Volume | $V$ |
| | XY Area | $A_{XY}$ |
| Graph | Graph density (Method 3.6) | $D$ |
| | Mean in- and out- degree and standard deviation (Method 3.6) | $d^{in}$ and $d^{out}$ |
| | Mean weight and standard deviation | $\bar{w}$ |
| | Mean Homogeneity (Method 3.6) | $\bar{H}$ |
| | Number of traversal steps (Method 3.6) | $\hat{k}$ |

### Topological heterogeneity revealed by the statistics of transition between neurons

Beyond size and density, we probed subnetwork topology via the mathematical theory of spectrum focusing on the properties of the transition probability matrices *P* (Fig. 6A; Methods 3.6) constructed from the connectivity matrix introduced above. Interestingly the second largest eigenvalue modulus (SLEM) of such matrix serves as a proxy for the mixing time (time for spiking in a single neuron to reach any other neuron inside the cluster) and reveals features such as bottlenecks in the network, well mixed organization (expansion quality), and homogenize connections between the neighbors of each neurons in the network (degree regularity) [33, 34, 35, 36]. Across all SCCs, SLEM values spanned a continuum (Fig. 6B), delineating two principal regimes:

**Figure 6:**
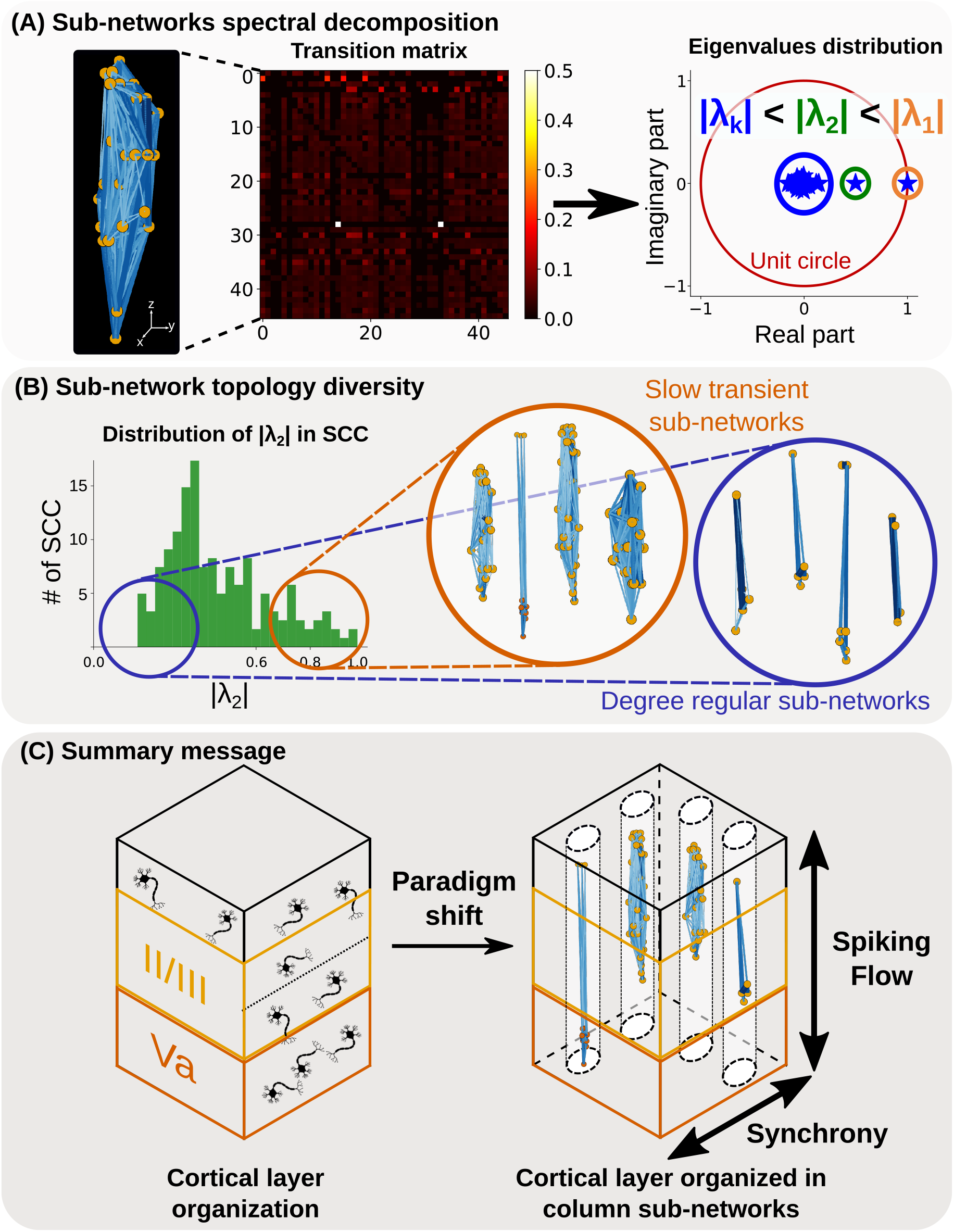
Spectral flow of neuronal activation across cortical sub-units. **A**. Left: Example of a sub-network representation with its normalized transition probability matrix *P*_*i,j*_ accounting for the transition between each neuron of the network. Right: Distribution of the eigenvalues *λ*_*k*_ of the matrix *P*_*i,j*_ represented on the complex plane. The first eigenvalue is 1 and the second (green) represents the speed of activation flow. **B**. Left: Distribution of second largest eigenvalue modulus |*λ*_2_| for all sub-networks. Right: Sub-networks topology trends for small (degree regular network [34, 36]) and large (slow transient networks or expander graphs [35]) |*λ*_2_|. **C**. Summary message : cortical layers can be decomposed into sub-networks organized in columns with a predominant feedback flow that can further fire in synchrony. These sub-networks could constitute computational sub-units.

1. **Low-SLEM, degree-regular modules**: characterized by rapid spike mixing and uniform link patterns, indicative of tightly knit, homogeneous clusters.
2. **High-SLEM, slow-mixing or expander-like modules**: featuring bottlenecks or hierarchical hubs that prolong information retention and support more complex routing.

Importantly, SLEM showed no clear dependence on SCC size, suggesting that both small and large neuronal subnetworks can manifest either topology. This spectral diversity underscores the repertoire of functional motifs embedded within the cortical columnar scaffold (see also Fig. S1).

## 2 Discussion

In this study, we combined fast volumetric two-photon calcium imaging with a novel graphbased inference pipeline to reconstruct three-dimensional functional connectivity in awake mouse primary motor cortex. From ∼1,000 neurons imaged over six 20-min sessions each recorded on different animals, we obtain a novel decomposition of cortical column network in sub-population derived from reconstructing weighted graphs and decomposed them into strongly connected components. Such decomposition allowed to uncover seven reproducible subnetwork archetypes—ranging from compact columns to diagonal and triangular motifs—predominantly in layer II/III but often bridging to layer Va. Quantitative analyses revealed a net feedback (Va→II/III) information bias, rapid access across nodes in ≤6 steps (with an average of 2), and both homogeneous and hub-dominated modules distinguished by their spectral profiles.

### Bridging simultaneous volumetric imaging and functional inference

Prior sequential two-photon studies in sensory areas provided key evidence for cortical columns via receptive-field mapping [7, 6], but their non-simultaneous sampling limits inference of temporal and causal interactions. In vitro calcium-imaging first demonstrated how optical signals could reveal functional microcircuits [37] and later how cortical layer II/III from primary visual cortex consists of an On and Off group of neurons leading to higher orientation selectivity and narrower bandwidth than individual neurons [38]. Finally, by applying our pipeline to true volumetric data recorded by sTeFo 2P *in vivo*, we now capture real-time, layer-spanning functional maps in M1, validating classic columnar motifs originally described in cats and monkeys [39, 40] while extending them to novel 3D geometries.

### Microcircuit motifs beyond the canonical column

In the barrel cortex, layer IV barrels show whisker-specific clustering, while those in layer II/III more diffuse tuning [41, 42], highlighting departures from strict radial modules. Our findings in M1 similarly reveal not only compact column-like SCCs but also elongated diagonal and triangular motifs. These diverse geometries echo anatomical observations of oblique intracortical projections [40] and point to a balance between vertical integration and lateral diversification in neocortical organization.

### Intrinsic versus task-evoked dynamics

Unlike task-driven mapping studies that combine encoding of kinematics or forces [43], our functional networks derive from ongoing, spontaneous activity—akin to the UP-state attractor dynamics originally characterized in slice preparations [37] or other manisfestation of spontaneous activity where ongoing cortical ensembles is similar to sensory evoked in groups of neurons with strong synaptic connectivity [44]. Overlaying task-evoked patterns onto network graph ensemble in future work should clarify how behavior recruits and reshapes specific subnetwork channels.

### Functional implications of subnetwork diversity and comparison with anatomical connectivity

The seven SCC archetypes exhibit distinct size, density, temporal profiles, and spectral signatures. The net feedback (Va→II/III) bias aligns with deep-to-superficial projections that integrate motor commands [45]. Homogeneous modules may serve rapid feedforward relay, whereas heterogeneous, hub-dominated clusters could implement flexible routing or gain control, similar to dense inhibitory subnetworks [46]. Low-SLEM (fast-mixing) motifs may support rapid sensorimotor loops, while high-SLEM (slow-mixing) modules facilitate temporal integration.

Anatomical connectivity has long been a gold-standard for neuronal circuit reconstruction, inferred from fixed-tissue reconstructions or transsynaptic tracing [47, 48, 49, 50]. These approaches provide static wiring diagrams but little insight into dynamic ensemble activity. While connectomic maps offer structural constraints [51], they themselves do not necessarily reveal which connections are functional *in vivo*, nor how neuronal ensembles encode stimuli over time [52]. Alternative approaches include functional circuit inference based on electrophysiology or fMRI data [53, 54]. Descriptive statistics, such as cross-correlogram, have been traditionally used to reveal underlying interactions between neurons, or information flow and hierarchy between brain areas [55, 56, 57]. As the number of recorded neurons increases [58], interpretation of such statistical structures becomes non-trivial [59, 60], even with the aid of anatomical model-based constraints [61]. More recent studies build on advances in calcium imaging and holographic optogenetics to infer and functionally validate cortical network structure from the dynamics of population activity. By selectively inactivating neurons and measuring the resulting effects on sensory encoding, it turns out that functional ensembles are not just subsets of anatomically connected cells, but are defined by their causal contribution to circuit computations, [38] and related work [44, 62, 63].

These efforts, complemented by algorithmic advances in spike inference and signal deconvolution [64], provide a comprehensive framework for mapping and interrogating dynamic network structure in behaving animals. Finally, our approach provides a dynamic, behaviorally relevant reconstruction of functional connectivity, enabling the identification of ensemble-specific roles that anatomical methods alone cannot resolve.

### Methodology limitations and future directions

Our adaptive, layer-specific thresholding provides high-confidence links but may omit weaker or transient interactions and underrepresent neurons projecting outside the imaged volume. The 4Hz sampling rate constrains temporal precision for very fast synaptic events, and deconvolution accuracy depends on calcium indicator’s kinetics. Combining this framework with higher-speed indicators or simultaneous electrophysiology could further refine connectivity estimates.

This generalizable pipeline could be applied to other cortical and subcortical regions, across developmental stages, or in disease models (e.g., Parkinson’s, stroke) to track microcircuit motif evolution. Integrating cell-type-specific perturbations will disentangle the roles of interneurons and projection neurons in each motif. Ultimately, correlating these subnetwork archetypes with behavior could elucidate elementary processing units of the cortex and their disruption in pathology.

## 3 Materials and Methods

### 3.1 Animals and ethics statement

This work complies with the European Communities Council Directive (2010/63/EU) to minimize animal pain and discomfort. Experimental procedures were approved by EMBL’s committee for animal welfare and institutional animal care and use (IACUC), under protocol number RP170001. 8-16 week old C57Bl6/j mice from the EMBL Heidelberg core colonies were used for experiments, housed in groups of 1-5 in makrolon type 2L cages on ventilated racks at room temperature and 50% humidity with a 12 hr light cycle. Food and water were available *ad libitum*.

### 3.2 Surgery

Surgeries were performed as detailed elsewhere ([65, 10, 11]). Briefly, 7-8 week old mice of either sex were used for cranial window surgeries. Animals were anesthetized with a mixture of 40 *µ*l fentanyl (1 mg/ml; Janssen), 160 *µ*l midazolam (5 mg/ml; Hameln), and 60 *µ*l medetomidin (1 mg/ml; Pfizer), dosed at 3.1 *µ*l/g body weight and injected intraperitoneally. After loss of pain reflexes, the fur over the scalp was removed with hair removal cream, eye ointment was applied (Bepanthen, Bayer) and 1% xylocaine (AstraZeneca) was injected under the scalp as pre-incisional local anesthesia. The mouse was then placed in a stereotaxic apparatus (David Kopf Instruments, model 963) equipped with a heating pad (37^°^C) to preserve body temperature. The dorsal cranium was exposed by removing the scalp and periosteum with fine forceps and scissors. Next, a 4 mm diameter circular craniectomy was made over M1 using a dental drill (Microtorque, Harvard Apparatus), centered at 1.75 mm anterior and 0.5 mm lateral to Bregma, avoiding damage to the dura and bleeding. For Ca^2+^ indicator expression at M1, recombinant AAV vectors (rAAVs, serotype 1) encoding jRGECO1a under the control of the synapsin promoter (Addgene #100854-AAV1) were stereotaxically delivered at the center of the craniectomy using glass pipettes lowered to depths of 300, 400, and 500 *µ*m, at a rate of ∼4*µ*l/hr using a syringe. Approximately 300 nl were injected per spot. After injection, the craniectomy was covered by a round 4 mm coverslip (∼170 *µ*m thick, disinfected with 70% ethanol) with a drop of saline between the glass and the dura. The cranial window was sealed with dental acrylic (Hager Werken Cyano Fast and Paladur acrylic powder) and a custom head fixation bar was also cemented. The surgical wound was closed with dental acrylic. Mice were single-housed after surgery to prevent compromising the implanted windows and had a recovery period of at least 4 weeks before imaging, allowing Ca^2+^ indicator expression and resolution of inflammation associated with surgery [66]. Post-surgical care included pain relief (Metacam, Boehringer Ingelheim) and antibiotic (Baytril, Bayer) subcutaneous injections (0.1 and 0.5 mg/ml respectively, dosed at 10 *µ*l/g body weight).

### 3.3 In vivo Ca^2+^-imaging of M1 populations

We used a self-built two-photon microscope based on scanned temporal focusing (sTeFo) enabling fast volumetric *in vivo* Ca^2+^ imaging of large M1 neuronal populations. The technical details and working principles are described in detail elsewhere [8]; however, for our study, we utilized a higher repetition laser (10 MHz, FemtoTrain, Spectra Physics) and operated the microscope with the following parameters: The scanning laser power was kept below 191 mW after the objective to avoid excessive heating of brain tissue [8]. Fields of view of 470 *µ*m^2^ were imaged at 128 px^2^ resolution and spaced in 15 *µ*m steps in the axial direction to cover 585 *µ*m in depth (39 z steps) at a volume rate of 3.99 Hz. Frames acquired during objective flyback were discarded. Typically, Ca^2+^ signals with good signal-to-noise were detected up to a depth of 500 *µ*m from the pial surface. Mice were briefly (*<* 1 min) anesthetized with 5% isoflurane in O_2_ for quick head fixation on a custom-built stage, with their bodies inside a 5 cm acrylic tube, under our custom two-photon microscope. After head fixation, mice fully recovered from the isoflurane anesthesia in under a minute, displaying clear blinking reflexes, whisking, sniffing behaviors, and normal body posture and movements. We waited at least 5 minutes before starting experiments to ensure full recovery from the brief anesthesia. Before Ca^2+^-imaging experiments in the awake condition, a pilot group of mice was habituated to the head-fixed condition in daily 20 min sessions for 3 days. We noted no marked behavioral difference between habituated and unhabituated mice during these relatively short 20 min imaging sessions. Imaging sessions typically lasted 20 minutes, after which the mouse was returned to its home cage. Typically, mice were imaged a total of four times (sessions), once per week. Among the four datasets, the one with the largest number of neurons detected (peak of AAV expression and cranial window quality) was considered for further analysis.

### 3.4 Functional neuronal network connectivity reconstruction based on calcium time series

We present here a step-by-step description of the computational approach we developed to transform 3D volumetric calcium neuronal data into 3D neuronal functional network (Fig. 1).

1. **Binarizing calcium time series:** we filter calcium responses after motion and neuropil correction using NoRMCorre and CaImAn [67, 28]. Deconvolve Ca^2+^ signal is binarized by extracting large peaks while excluding low-amplitude peaks through thresholding (Fig. 1A).
2. **Removing clustered population spiking based on background statistics:** to avoid counting population responses, we introduce a mapping function Φ that randomly removes spikes present in population spiking intervals using the non-bursting time intervals statistics (Fig. 1B).
3. **Identifying time correlation between neurons:** we develop a sequential analysis on all calcium time series to define the successive activation and co-activation events between neurons. The result is a temporal correlation measure for each pair of neurons *i, j* based on their sequential and synchronous relationship (Fig. 1C).
4. **Graph reconstruction of the network:** the neural network is constructed by creating a graph with an adjacency matrix *A*_*i,j*_. Each element *A*_*i,j*_ corresponds to the functional temporal correlation measure and represents the strength of the connection between neurons *i* and *j* (Fig. 1D).
5. **Removing sparse correlation by thresholding the correlation matrix:** to eliminate spurious connections, we created a statistical null-hypothesis informed by dataset-specific parameters. This model defines a maximum coincidental weight for random connections used as a threshold to distinguish genuine connections from spurious ones (Fig. 1D).
6. **Neuronal network reconstruction using an adaptive threshold for each layer:** to account for variations in firing rate distributions and inter-layer connectivity dynamics, we threshold the graph depending on each neuronal layer statistics (Fig. 1E).
7. **Graph partition into high-connectivity sub-networks:** to identify highly correlated sub-networks, we decompose the graph of the network into strongly connected components (SCCs), see Fig. 3A. Outlier sub-networks are identified using a machine learning approach and re-decomposed based on the connections statistics (Fig. 3B). Finally, smaller SCCs are excluded to retain only functionally relevant clusters of activity.
8. **Construction of coarse-grained network:** we extract binary activation signals from SCCs and construct a coarse-grained network using Spike Time Tiling Coefficient [31] of pairs of signals as connection weight between sub-networks (Fig. 3D).
9. **Classification of sub-networks in subtypes:** after extraction of relevant biological, geometrical and functional features using PCA, SCCs are classified in seven types based on their characteristics using machine learning (Fig. 5A).

We now describe the details of each step.

#### 3.4.1 Ca^2+^-imaging signal extraction

Volumetric imaging data were visualized and rendered using FIJI [68]. Motion correction of *in vivo* awake recordings was performed using NoRMCorre [67] on each imaging plane in the volumetric datasets. jRGECO1a signal was extracted, filtered, and neuropil-corrected from each individual imaging plane using CaImAn [28]. For each neuron, its corresponding deconvoluted time-series is processed by detecting its peaks using the find peaks function of the *scipy* library with prominence *p* = 0.015. A spike is detected when the peak reaches the threshold *T*_*f*_ = 0.05 (see Table 3). The peaks are used to binarize the signal and create a spike train where the neurons is either activated or not (Fig. 1A).

**Table 3:** Parameters used for network reconstruction and segmentation.

| Parameters | Symbol | Values |
| --- | --- | --- |
| Peak detection prominence | $p$ | 0.015 |
| Spike firing threshold | $T_f$ | 0.05 |
| Activation regime threshold | $T_c$ | $2\sigma$ |
| Adaptive threshold confidence | $\alpha$ | 0.05 |
| SCC Size threshold | $T_{scc}$ | 3 |
| XY plan distance threshold | $T_D$ | 45 |
| Event activation threshold of an SCC | $T_{event}$ | 2 |

#### 3.4.2 Time series pre-processing based on background statistics

Despite the head-fixed condition of the mice, three out of six datasets contained recordings of clustered spike events of neurons corresponding to different regimes of activation. These regimes are characterized by different mean activity rates, which, with our network reconstruction approach, are more likely to introduce noise rather than convey informative data. However, we believe relevant clues about the underlying network structure can be retrieved from those regimes. Rather than discarding these events [13], we propose the following denoising method that preserves the continuity of neuron activation across different regimes. Random clustered spike events are detected from the mean spike train by using the threshold *T*_*c*_ = 2*σ*, with *σ* as the standard deviation of the signal (see Table 3). This approach segregates the time series into two distinct set of intervals: the background regimes and the clustered spike events, each characterized by a different mean firing rate 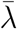.To align the clustered spike intervals with the background regimes, we introduce a mapping function *ϕ*. For each neuron *i, ϕ* adjusts the clustered spike intervals by randomly removing spikes until the interval matches the firing rate of the surrounding background regime (Fig. 1B & Fig. S2). This non-parametric approach is resilient to variation of the background activation and preserves continuity within the signal of a neuron independently of others. This continuity is a necessary hypothesis to define the thresholding method explained later on (Method 3.4.5).

#### 3.4.3 Identifying time correlation between neurons

Due to the fast firing rate of neurons compared the limited image acquisition frequency of the calcium signaling, genuine time ordering of spikes may not be feasible when they occur in the same time bin. To address this limitation, the following approach is suggested (Fig. 1C). We define the strength of the connection between two neurons *i* and *j* as the weight *w*_*i,j*_. When *w*_*i,j*_ is null, no connections exists between *i* and *j*.

1. **Successive correlated events**. When two spikes are occurring one after the other, a single directed edge is created from *i* to *j* (Fig. 1C upper), leading in an increment of *w*_*i,j*_ by one.
2. **Synchronous correlated events**. In instances where two spikes are occurring simultaneously, no temporal order can be determined within this *same bin interaction* [13]. Consequently, two directional edges are added between neurons *i* and *j* : one from *i* to *j* and one from *j* to *i* (Fig. 1C - center). This results in an increase of *w*_*i,j*_ and *w*_*j,i*_ by one.

Considering the spatial scale of the data (∼500*µm*^3^), no spatial distance can be determined to influence the causal relationship between subsequent firing neurons. Meaning no choice can be made to preferably connect one neuron to another. Moreover, self-connections are excluded, ensuring no neuron forms a connection with itself. This strategy makes sure that the real functional path is included in the set of all created paths. Given the long duration of the recordings (∼20 minutes), we assumed that true activation pathways would reoccur consistently over time. By a thresholding step, only the repeatedly occurring (i.e. strongly correlated) connections will be kept.

#### 3.4.4 Reconstructing the neural network

A directed weighted graph *G* is generated to represent the neural network. The graph is associated with an adjacency matrix *A* of dimension *N* ^2^ with *N*, the number of neurons. First, the weight between every pair of neurons *i* and *j* is computed by applying an iterative algorithm using the previously mentioned principles over the total recording duration *T* :

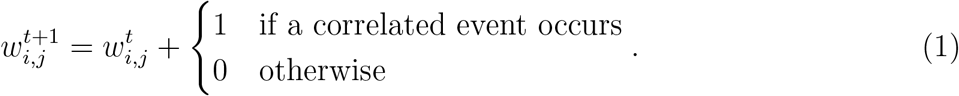

Then, the final weight 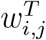 is stored in each corresponding *A*_*i,j*_. In order to simplify the calculation, the graph can be computed using the spike trains representation as a set of activation times *s* = {*s*_1_, *s*_2_, …, *s*_*n*_}. Then, our expression of *A*_*i,j*_ for a directed network is determined by the spike trains *s*_*i*_, *s*_*j*_ of neurons *i* and *j* and a time shift *δT* = 1:

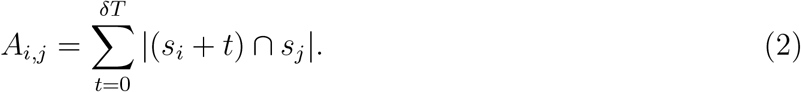

#### 3.4.5 Threshold computation criteria

The resulting adjacent matrix *A*_*ij*_ from the previous step contains numerous irrelevant coincidental connections characterized by small weights. We decided to create a deterministic adaptive threshold to remove these spurious connections and thus manage to keep only the ones with the highest probability to be genuine functional pathways (Fig. 1D). However, defining such threshold is not trivial [69]. We propose a threshold estimation approach taking into account the dataset characteristics and the difference among neuronal layer populations. This approach computes the maximal coincidental connection that appears from independent Poisson processes, used as a statistical null-hypothesis. Each neuron *i* in a population *L* is characterized by a firing rate *λ*_*i*_. If all neurons in *L* follows an independent Poisson process, then for any two neurons *i, j* ∈ *L*^2^, the probability that both neurons fire synchronously or consecutively within Δ*T*, is given by:

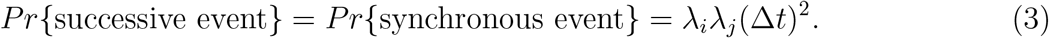

By summing these probabilities, the probability of increasing the weight connection between *i* and *j* in our graph construction is:

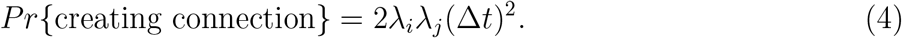

Given the small magnitude of this probability, the number of connections can be approximated by a Poisson distribution with parameter 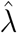, which depends on the maximal connection probability and the number of samples. To determine the maximal coincident connection probability, we consider the two most active (i.e. highest firing rates) neurons inside the population *L*, characterized by firing rates *λ*_*m*_, *λ*_*m*−1_. The number of samples 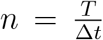 is calculated using the total recording duration *T* and Δ*t* = 1:

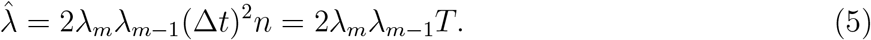

Thus the probability of observing *k* or less connections between the most active neurons is computed from the cumulative distribution function (CDF) of 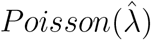 distribution:

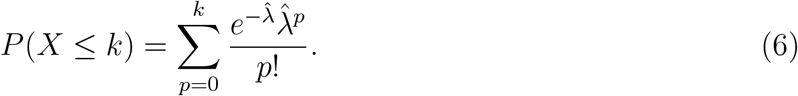

Using the CDF, we calculate the smallest *k* for which *P* (*X* ≥ *k*) *< α*, with the confidence parameter *α* = 0.05 (see Table 3). This approach ensures the choice of threshold *k* to filter out random connections and can dynamically adapt the threshold to data-specific properties by varying the chosen population and the number of samples. However, this method assumes that the number of samples is sufficient and the firing rates small and stationary.

#### 3.4.6 Adapting the threshold to layer dynamics

The thresholding procedure removes invalid connections appearing fewer than *k* times in a population *L*. The threshold acts as a maximum coincidence boundary, filtering out spurious connections and potentially some real connections, while ensuring that the retained connections are statistically robust and unlikely to occur by chance. In our datasets, this threshold is not uniform but varies across layers and between them, reflecting differences in firing rate distributions and connectivity among neuronal layer populations (Fig. 1E & Fig. 2A). For inter-layer connections, the threshold is derived from the maximal firing rates of the respective populations. By employing a layer adaptive null-hypothesis with a fixed data-independent parameter *α*, this approach avoid the need for manually defining a hard threshold. However, this strict filtering process also results in some neurons being excluded from the active network, as their activity lacks sufficient correlation with other neurons within the region to meet the thresholding criteria (Fig. 2A).

#### 3.4.7 Random networks as a statistical comparison

To compare network quantities when the threshold is increased (see Figs. S3-S5), we simulated random networks where each neurons follows an homogeneous Poisson point process with their average firing rate as parameter. For each dataset, 50 simulations were computed of equal duration as the recordings. A measure of convergence *r*_*scc*_ is established to compare the emergence of sub-networks within the networks (see Fig. S5) and is computed with the number of sub-network edges *card*(*E*_*scc*_) (see Methods 3.5.1) and the total number of edges *card*(*E*) :

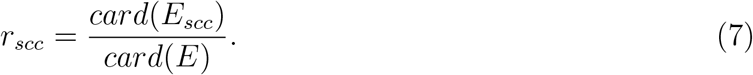

### 3.5 Sub-network decomposition, classification and activity patterns

#### 3.5.1 High-connectivity sub-networks obtained from graph decomposition

In order to find relevant sub-networks within the created graph, a strongly connected component (SCC) partitioning is performed using Pearce algorithm [70] (Fig. 3A). A directed graph *G* = (*V, E*) is called strongly connected if for every vertices *i, j* ∈ *V* ^2^ there is a directed path from *i* to *j*. The strongly connected components of a directed graph are a partition into sub-graphs each of which is strongly connected (Fig. 3A). From a Markovian perspective, we are dividing the graph into irreducible Markov chains. Each SCC represents a set of neurons that seems to have similar pattern of activity in space and time. After applying the algorithm, we found out that most of the SCC had a column shape except for a few outliers. Those outliers were defined by several columnar shaped clusters strongly connected together. For statistical comparison purposes, we decided to decompose these outliers into elementary columns that remains strongly connected. The approach is the following :

1. **Outlier detection**. A DBSCAN algorithm [30] was employed to automatically find these outliers among SCCs using the number of neurons *N*_*c*_ and the spread *S*_*scc*_ on the XY plane projection (see Eq. 8) as discriminative features (Fig. 3B).
2. **Separation in columns**. Lengthy connections between neurons were removed based on their pairwise XY-plane distances to separate the cluster into columns. The distance threshold, denoted as *T*_*D*_ = 45, was empirically set based on the distribution of distances between neurons in the outliers (see Fig. 3B, Table 3).

This two-step process allowed us to dissect the large synchronous components into elementary substructures comparable to other SCCs, facilitating a more detailed characterization of the network’s functional organization. Only sub-networks with a number of neurons *N*_*c*_ *> T*_*scc*_ = 3 (see Table 3) were kept after applying our method (Fig. 3A).

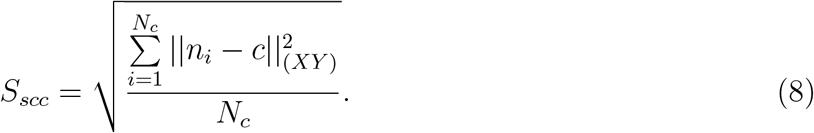

#### 3.5.2 Coarse-grained network using sub-networks as elementary components

An event within a strongly connected component (SCC) is defined as a continuous sequence of activations involving a minimum of *T*_*event*_ = 2 (see Table 3) within the sub-network. An event concludes when no further neuronal activation occurs following the last activation. With this definition, we extract a signal of activation of each SCC. Using this signal, we create a pairwise functional matrix representing a coarse-grained network between SCCs (Fig. 3E). This matrix is created using the Spike Time Tiling Coefficient (STTC) [31], a measure of correlation usually between pairs of neurons that is independent of firing rate. In order to fit in the STTC computation, we only use the start of the events to describe the SCC spike train. The time window is empirically set to four time bins (∼ 1s) because most the events in the SCC are a second long. A coarse-grained graph that represents the synchrony between SCC is thus obtained using the pairwise functional matrix.

#### 3.5.3 Subtypes of cortical layer sub-networks revealed by K-means classification

After the partitioning approach was applied, strongly connected components (SCCs) were classified into several subtypes based on both geometrical and functional properties. First, we use the biological layer information to classify SCCs into three types: SCCs with nodes exclusively in Layer Va, exclusively in layer II/III, and SCCs in both layers. However, as illustrated in Fig. 3C, the distribution of these classes was found to be highly imbalanced, with the majority of SCCs originating from layer II/III. To refine the classification further for layer II/III clusters, we define a vector of 27 features that are listed below in Table 2. We use a principal component analysis (PCA) to reduce the dimensionality from 27 to 6 dimensions. The number of output features was selected to retain between 70% and 80% of the explained variance ratio. The K-Means clustering algorithm was applied on the layer II/III SCCs feature vector to identify more specific subtypes. The K-Means optimal number of clusters was determined using the elbow method, resulting in the identification of five distinct subtypes of columnar SCCs within Layer II/III (Fig. 5A).

#### 3.5.4 Events activation within sub-networks

SCCs activation events (see Methods 3.5.2) also enables the characterization of activation regimes associated with SCC subtypes by computing event statistics for each class, as presented in Fig. 5B. The number of activations can be quantified for nodes serving as the first or last in an event sequence. This approach facilitates the comparison of the normalized relative depths of the starting and ending nodes of events, as illustrated in Fig. 4H.

### 3.6 Network statistics

In this section, we define the statistical quantities we used here. The neuronal network is represented by directed weighted graph *G* associated with an adjacency matrix 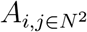 [71, 15], made of *N* neurons.

1. **The degree** is the number of in and out neurons given respectively by

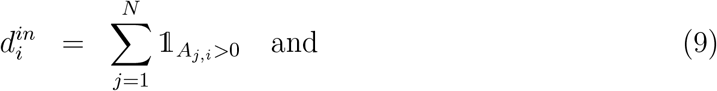

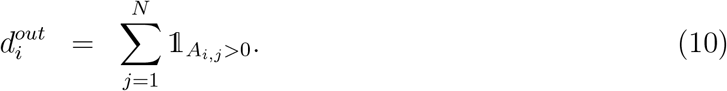
2. **Connection density** is the number of connections over the total number. It can be computed using the adjacency matrix as

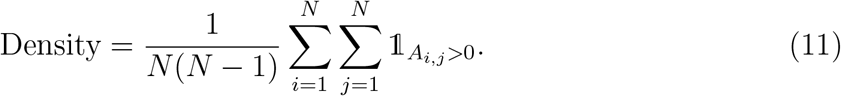
3. **Asymmetry** *As* is the difference in connection strength between two layers. For a neuron *i* in layer *L*_1_ and *j* in layer *L*_2_, we have

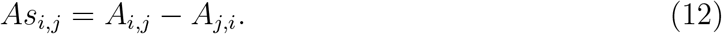
4. **Homogeneity** represents the preferential connections to some particular nodes compared to other. It is defined as the fraction between the maximal and the minimal outward connection strengths for a given node *i* :

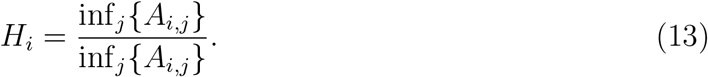
5. **Graph traversal number of steps** 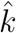 is the minimal number of iteration to transform the transition probability matrix *P* into a strictly positive matrix. We recall the adjacency matrix can be normalized into a transition matrix of probability *P* to represent the graph as a Markov Chain [32]. This number describe the accessibility of each neurons in the graph defined as

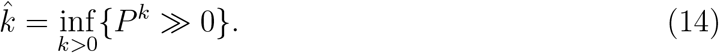

The symbol ≫ means that all coefficient of the matrix are stricly positive.
6. **Second largest eigenvalue modulus (SLEM)**: *λ* = {*λ*_1_, *λ*_2_, …, *λ*_*k*_} is the set of eigenvalues of the previously defined transition probability matrix *P* such that 1 = *λ*_1_ *>* |*λ*_2_| ≥ |*λ*_3_|≥ … ≥| *λ*_*k*_|. The modulus of *λ*_2_ is proportional to the convergence rate of the Markov Chain defined by *P* [33] and is a proxy of the topology of the underlying graph (see Fig. S1).

## Acknowledgments

DH is supported by ANR AstroXcite, Memolife and the European Research Council (ERC) under the European Union’s Horizon 2020 research and innovation program (No 882673). NR is supported by ANR (AstroXcite) and Memolife. JCB acknowledges supporting fellowships from the EMBL Interdisciplinary Postdoc (EIPOD) Programme under Marie Sk-lodowska Curie Cofund Actions MSCA-COFUND-FP (664726). JCB and RP were funded by the European Molecular Biology Laboratory (EMBL) and the Deutsche Forschungsgemeinschaft (DFG, German Research Foundation) – projects 458898724 and 425902099) warded to RP.

## Supplementary Materials

**Figure S1:**
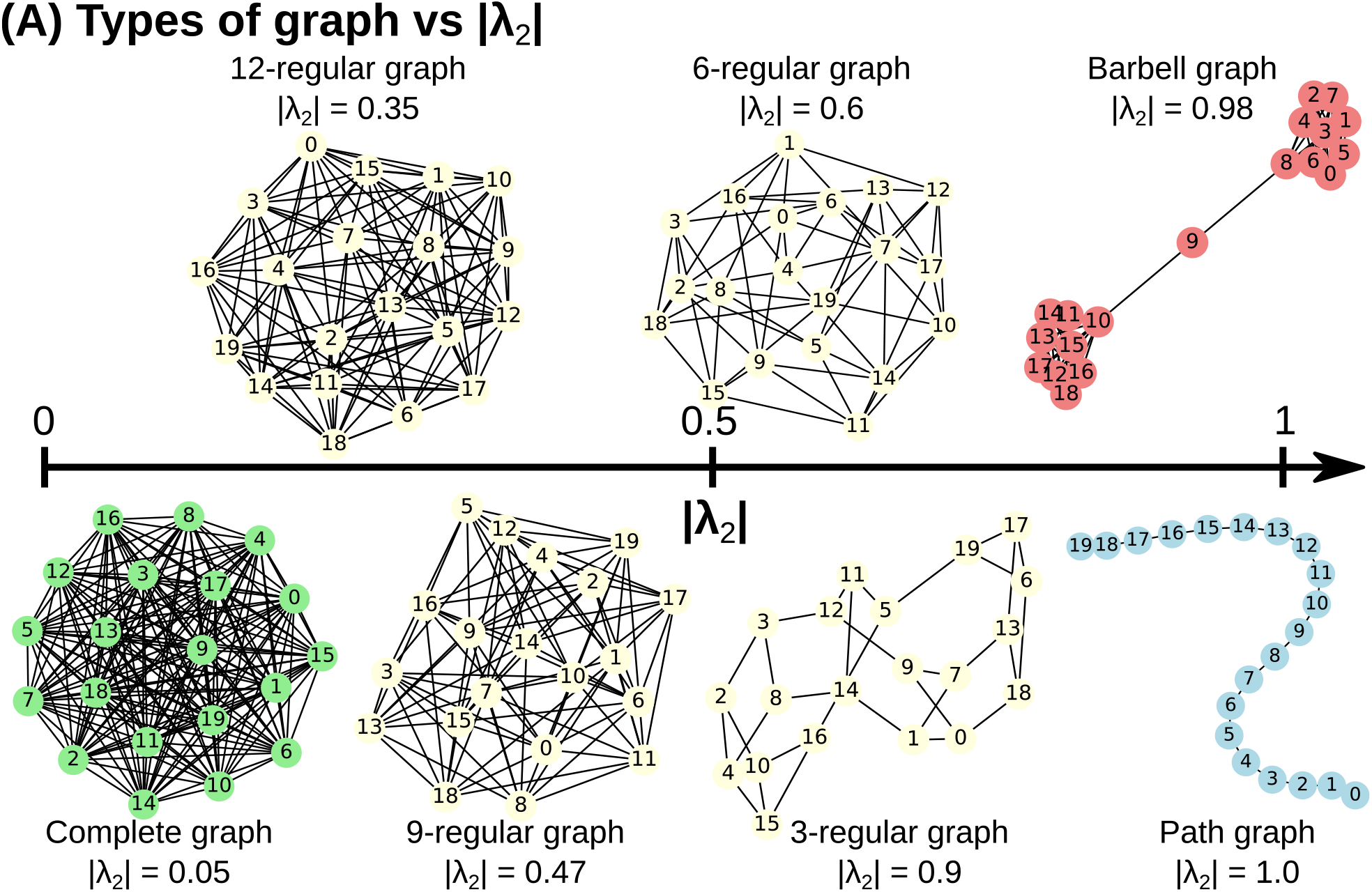
Types of network topology vs eigenvalues. **A**. From left to right, different graph representation as we increase the second largest eigenvalue modulus |*λ*_2_| (see Methods 3.6)

**Figure S2:**
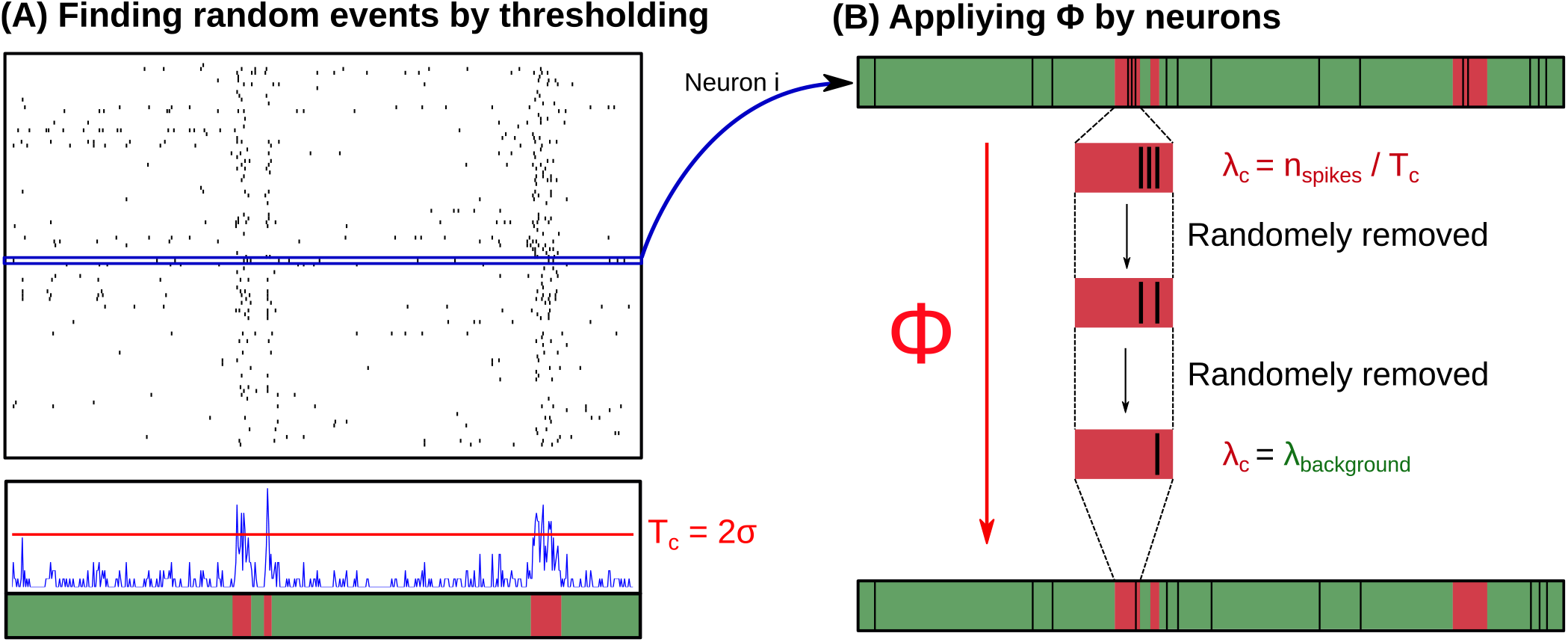
Function Phi illustration. **A**. Computation of the sum of all spike trains (blue) and extraction of clustered spike events by applying threshold *T*_*c*_ *>* 2*σ* **B**. For each neurons, function *ϕ* is iteratively randomly removing spikes in clustered spike events intervals (red) until intervals matches the background firing rate (green).

**Figure S3:**
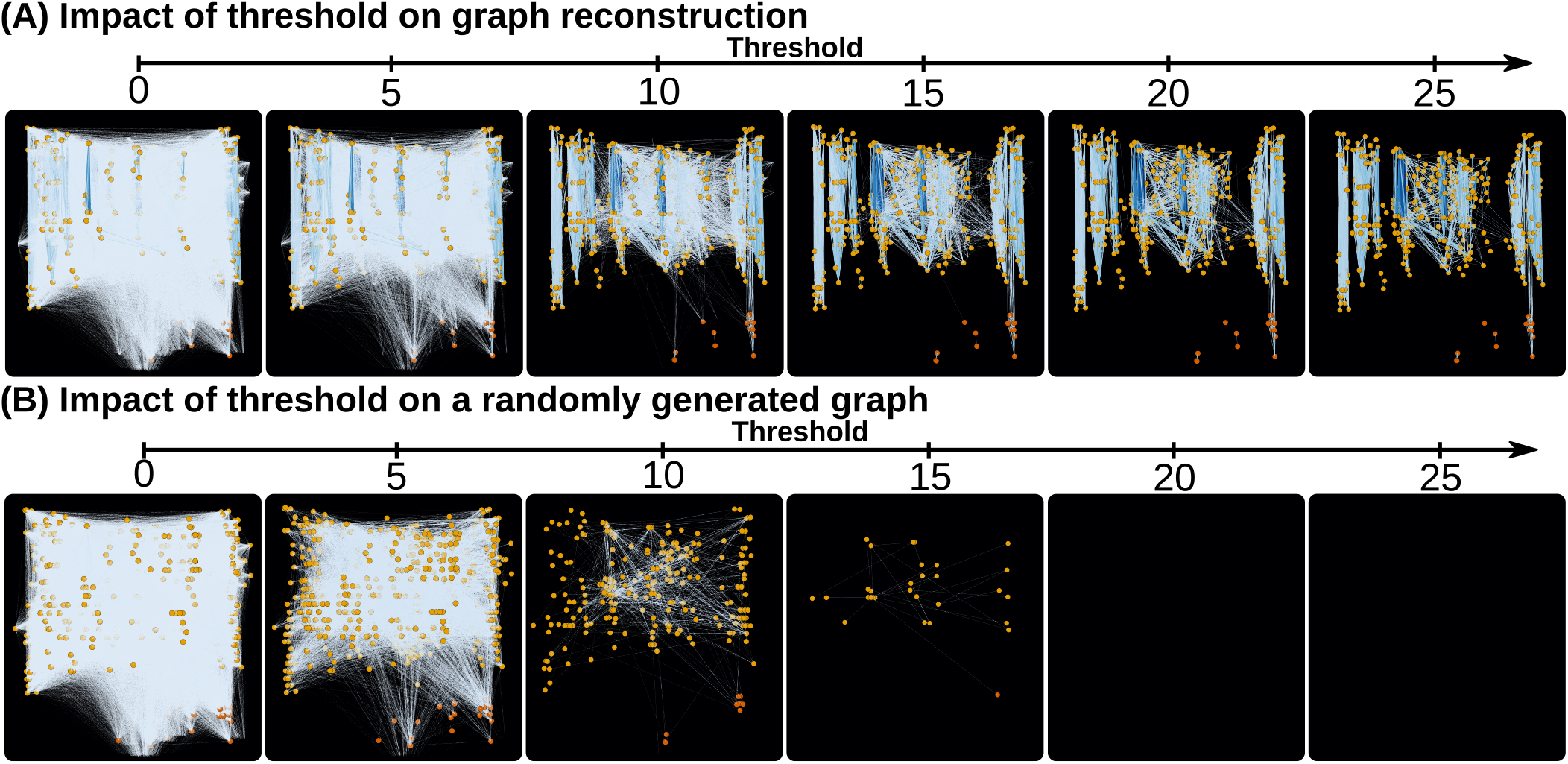
Impact of threshold on graph reconstruction. **A**. Impact of the threshold’s increase on the network. **B**. Impact of the threshold’s increase on independent Poisson process random network.

**Figure S4:**
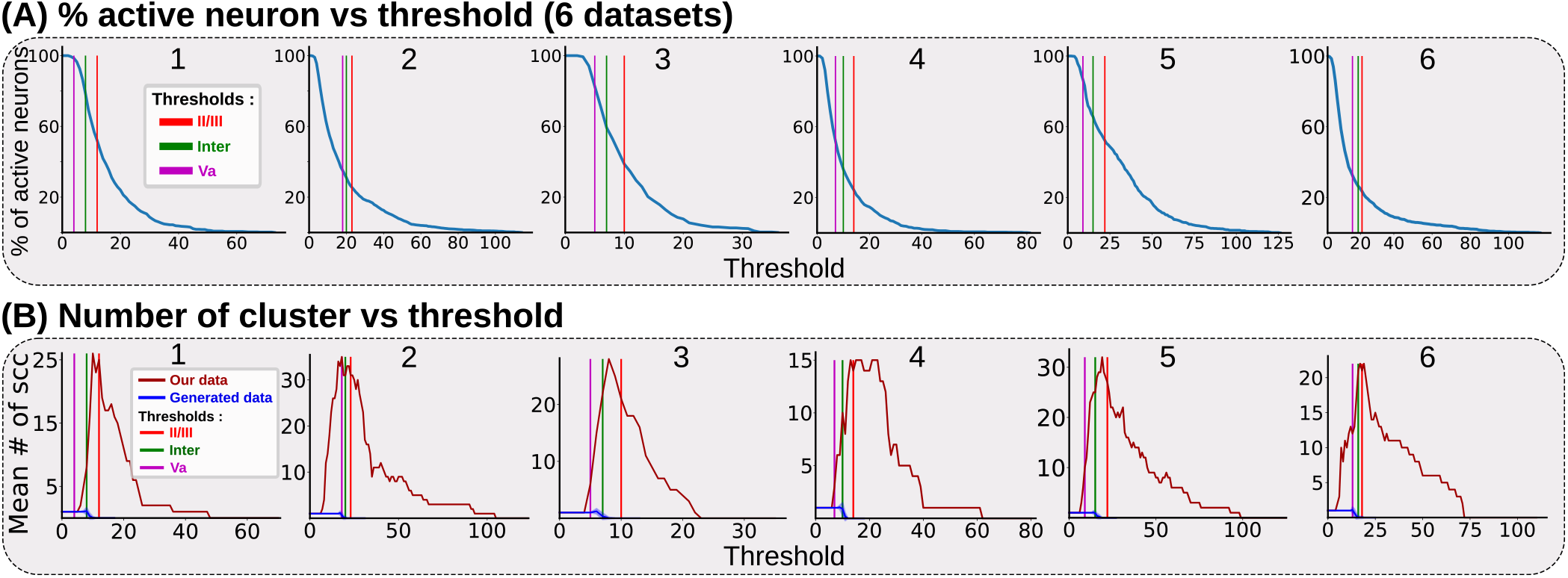
Network statistics vs threshold. **A**. Proportion of active neurons with at least one connection in the network when increasing threshold *T* for each 6 experiments. **B**. Number of SCC when the threshold increases for each 6 experiments (red) and for randomly generated networks (blue).

**Figure S5:**
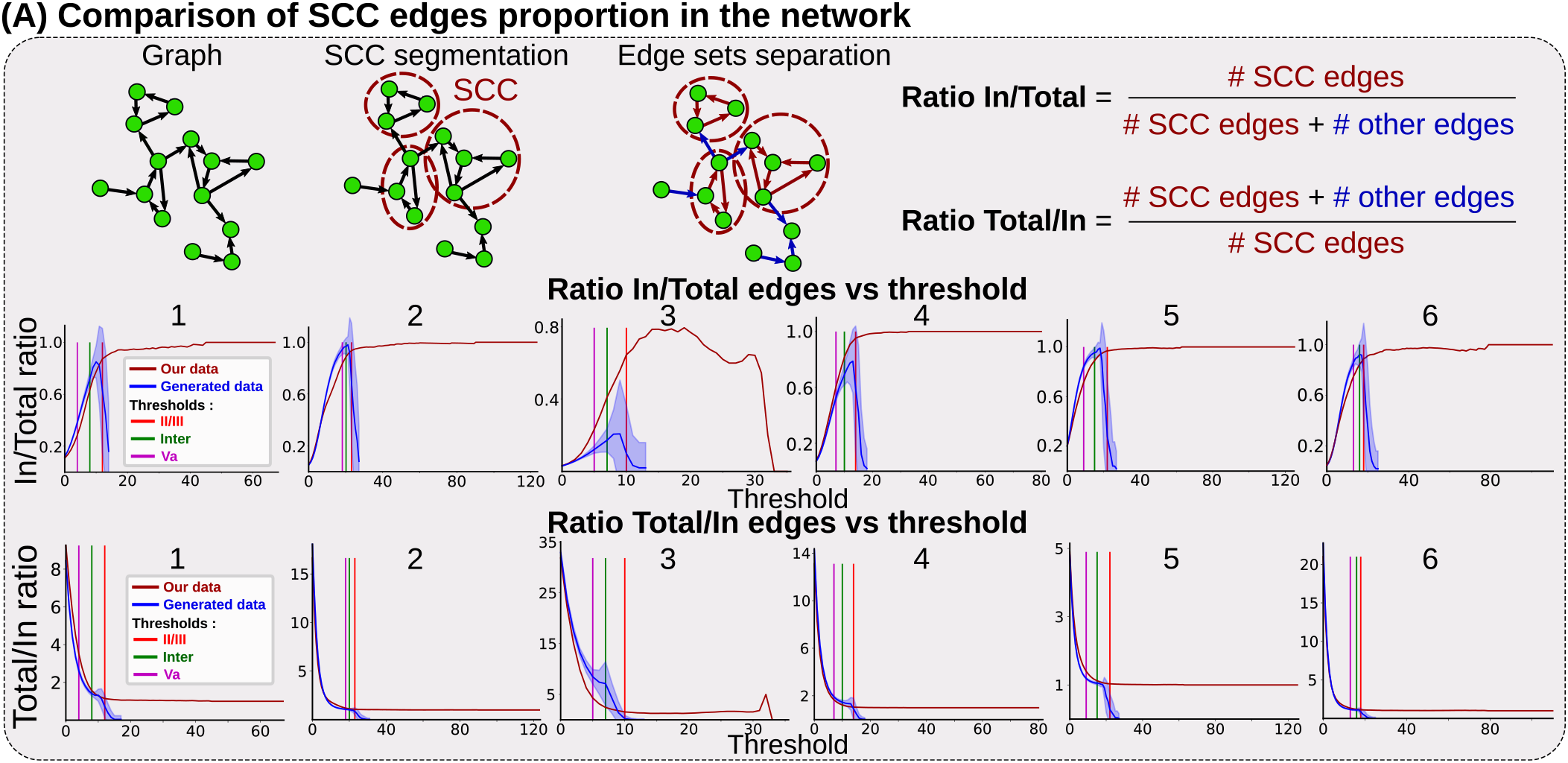
Clustered edge ratio vs threshold. **A**. For each experiments, SCC are computed and the ratio between the number of edges in the sub-networks vs the total number of edges is computed for each threshold *T* as well as the inverse ratio (red). The two ratios are computed for randomly generated data (blue).

